# Volatile aroma compound production is affected by growth rate in *S. cerevisiae*

**DOI:** 10.1101/2022.09.01.506294

**Authors:** Federico Visinoni, Penghan Zhang, Katherine A. Hollywood, Silvia Carlin, Urska Vrhovsek, James Winterburn, Daniela Delneri

## Abstract

The initial growth rate of a yeast strain is a key parameter in the production of fermented beverages. Fast growth is linked with higher fermentative capacity and results in less slow and stuck fermentations unable to reach the expected final gravity. As concentrations of metabolites are in constant state of flux, quantitative data on how growth rate affects the production of aromatic compounds becomes an important factor for brewers. Chemostats allow to set and keep a specific dilution rate throughout the fermentation and are ideal system to study the effect of growth on aroma production. In this study, we run chemostats alongside batch and fed-batch cultures, compared volatile profiles detected at different growth rates, and identified those affected by the different feeding profiles. Specifically, we quantified six abundant aroma compounds produced in anaerobic glucose-limited continuous cultivations of *S. cerevisiae* at different dilution rates. We found that volatile production was affected by the growth rate in four out of six compounds assayed, with higher alcohols and esters following opposite trends. Batch and fed-batch fermentations were devised to study the extent by which the final concentration of volatile compounds is influenced by glucose availability. When compared to the batch system, fed-batch fermentations, where the yeast growth was artificially limited by a slow constant release of nutrients in the media, resulted in a significant increase in concentration of higher alcohols, mirroring the results obtained in continuous fermentations. This study paves the way to further process development optimization for the production of fermented beverages.

**Importance:** The production of fermentation beverages will need to quickly adapt to changes in both the climate and in customer demands, requiring the development of new strains and processes. Breakthroughs in the field are hindered by the limited knowledge on the physiological role of aroma compounds production in yeast. No quantitative data on how growth rate affects aroma profile is available in the literature to guide optimisation of the complex flavours in fermented beverages. In this study, we exploited the chemostat system, alongside with batch and fed-batch cultures to compare volatile profiles at different growth rates. We identified the aromatic compounds affected by the different feeding profiles and nutrient limitations. Moreover, we uncovered the correlation between yeast growth, esters and higher alcohols production. This study showcases the potential of the application of feeding profiles for the manipulation of aroma in the craft beverage industry.

## Introduction

The invention of brewing, as fermentation of cereal sources to produce alcoholic beverages, pre-dates history and represents a hallmark of civilization (1). Brewing yeast strains underwent centuries of domestication, being improved for growth and aroma profile through spontaneous mutations and selection. However, while the scientific literature on the brewer’s yeast, *S. cerevisiae*, is extensive and detailed, little is known of the physiological role of the most abundant aroma compounds nor of the biosynthetic mechanisms which regulate their production. Amongst the broad and diverse range of compounds responsible for the complex aroma and taste of fermented beverages, the main yeast-derived contributors to the flavour of aroma and beer are higher alcohols and esters (2). These compounds give a pleasant, often fruity aroma to the beer when present in moderate quantities and are closely connected to the yeast’s main metabolism, being produced as secondary metabolites in trace quantities (3).

Higher alcohols are produced as a result of amino acid metabolism in the Ehrlich pathway through the catabolism of amino acids or through an anabolic route from pyruvate (4). Esters are produced by a condensation reaction between an alcohol and acetyl-CoA (acetate esters), or between ethanol and acyl-CoA (fatty acid esters) (5). Due to the high concentrations of higher alcohols reached in the fermentation process and the low odour threshold level of esters, a lot of effort has gone into improving their production through synthetic biology approaches (6–8). However, to successfully apply adaptive evolution approaches and fine-tune their production, the physiological role of the synthesis of higher alcohols and esters must be understood.

Researchers must also face the complexity of the fermentation process, as brewing is generally conducted as a batch process where both the fermentation parameters and the concentrations of metabolites are in constant state of flux (9). The process is also carried out without agitation and in conditions of transient aerobiosis as the fermentation starts with a pulse of oxygen which favours the production of unsaturated fatty acids and sterols (10). For these reasons, the fermentation step is characterized by an accentuated unsteady state, in which yeast physiology cannot be followed nor easily described (9, 11). Nonetheless, to develop new rational approaches for the optimization of fermentation processes it is necessary to understand how the ever-changing cell physiological state may affect and regulate production of aroma compounds.

Thus, it is of foremost importance to follow the fermentation process in a controlled environment where the agents contributing to shaping the aroma profile can be singled out and studied. A key tool in achieving this is the chemostat, a bioreactor in which cells grow continuously in a fixed volume, thanks to continuous addition of medium and removal of an equivalent volume culture. In fact, eventually, these systems are able to reach a condition of equilibrium, or steady state, in which the growth rate of the cells and the dilution rate are equal and constant and metabolite concentrations are stable over time (12, 13).

Several studies have been carried out to describe the metabolic fluxes of *S. cerevisiae* strains in anaerobic, glucose limited continuous cultures at different dilution rates (14, 15). However, the scope of these studies was limited to the characterization of yeast core metabolism and not of the secondary pathways responsible for the production of volatile compounds. More recently, classic continuous fermentations have been conducted to study the effect of oxygenation, temperature and nitrogen sources on the aroma profile (16–18), though, no quantitative data on how specific growth rate affects aroma profile during fermentations is currently available in the literature.

In this study, the link between yeast growth rate and aroma compound production is elucidated by characterizing the production of the most abundant higher alcohols and esters in continuous culture at different dilution rates in a *S. cerevisiae* type strain (NCYC 505). The results obtained from the chemostat cultures were subsequently used to devise feeding strategies to modulate the production of both higher alcohols and esters (Figure 1).

**Figure 1.**
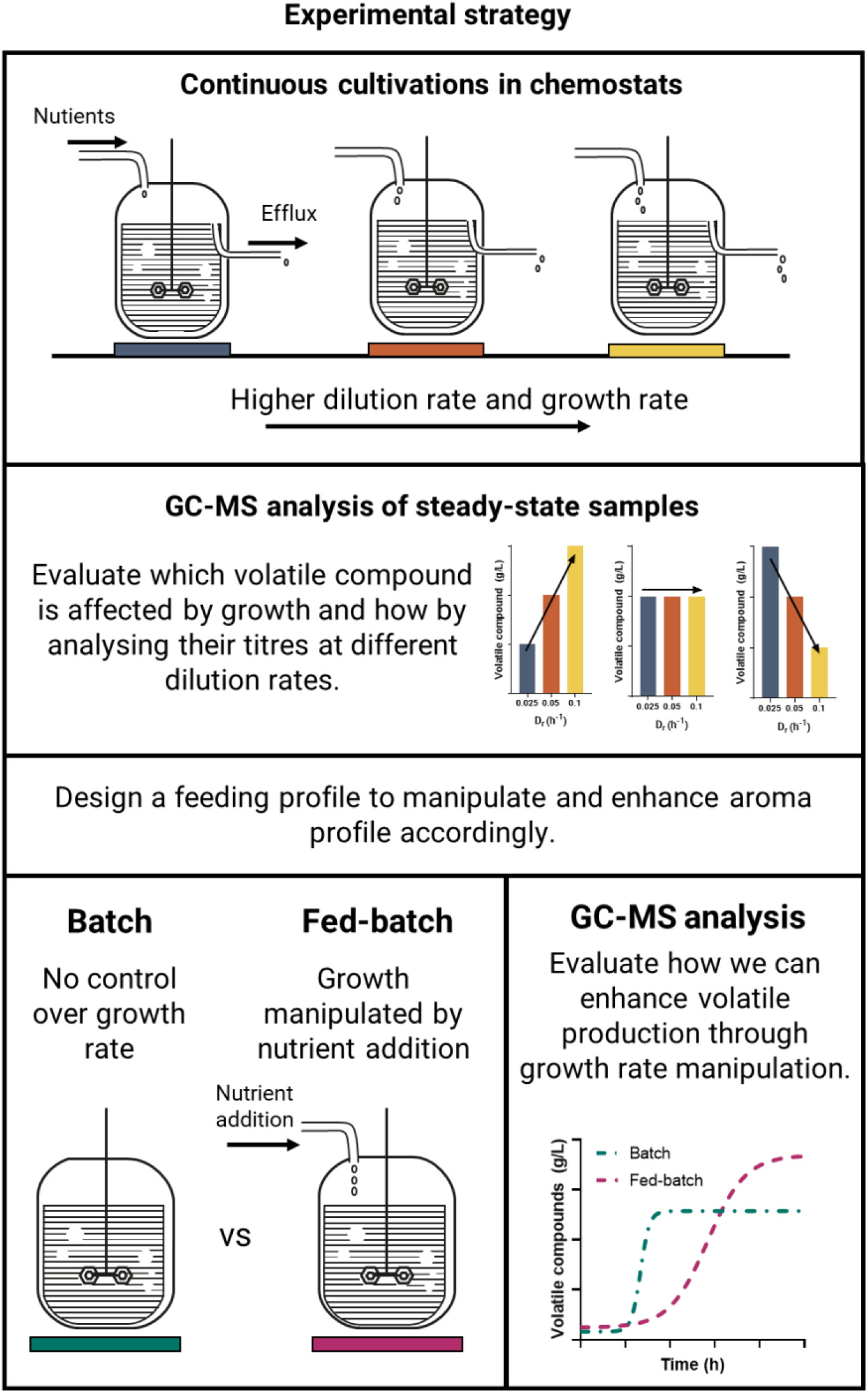
Experimental strategy. Continuous cultivations in chemostats will be employed to single out the effect yeast growth rate could have on the production of volatile compounds. The resulting GC-MS data from steady state samples at different dilution rates will be used to design a feeding profile to enhance the production of selected aroma compounds by artificially manipulating growth rate through nutrients additions.

Our study show that the production of volatile compounds can be enhanced and manipulated to yield different and improved aroma profiles via the implementation of feeding profiles. Moreover, these results will help defining synthetic and experimental evolution approaches to generate strains able to produce higher titres of desired volatile compounds in both wine and beer fermentations.

## Results

### Metabolic characterization of NCYC 505 in continuous carbon-limited cultures

To single out the effect exerted by the growth rate on the production of aroma compounds continuous cultivations in glucose limited chemostats were performed. The cultures were initially grown in batch mode, until 50 hours when the continuous culture was activated with a constant feed of YNB + 2% glucose at different dilution rates (D_r_): 0.025, 0.05, and 0.1 h^−1^

We found that the biomass decreased linearly with dilution rate, an effect likely due the uncoupling between anabolism and catabolism under substrate sufficient conditions (15, 19). Efflux samples at the steady state for the cultures grown at 0.025 and 0.05 h^−1^ D_r_ showed little to no glucose in the medium, whilst at 0.1 h^−1^ D_r_, 7.1 g/L of residual glucose was detected. Nonetheless, no washout or decrease in biomass was registered during the 200 hours of fermentation. The main exometabolites were first analysed using HPLC. Ethanol concentrations decreased with increasing growth rate, while its productivity and specific concentration (calculated as g/(L·OD_600_)) increased at higher dilution rates as a result of the increased availability of carbon source (Table 1, Figure 2). Similarly, a positive correlation between growth rate and glycerol production was observed, which can be easily attributed to its vital role in balancing the NADH used in anabolic reactions in anaerobic conditions (14, 15). Moreover, at the highest dilution rate tested, a significant increase in the concentration of acetic acid was recorded. Such production was at a higher rate than previously described for other yeast strains (14) (Figure. 2). Given that residual glucose was detected in the medium at 0.1 h^−1^ D_r_, it is likely that the acetic acid production is due to the overflow metabolism, as seen in other species (20–22).

**Table 1.** Biomass, specific and volumetric concentrations of exometabolites and yield on substrate from chemostat experiment with NYCY 505 in YNB + 2% glucose at different dilution rates. Yield is represented by the ratio between Qp, specific productivity (metabolite concentration (g/L) ∙ flow rate (L/h) / volume (L)), and Qs, specific substrate uptake rate (substrate feed concentration (g/L) ∙ flow rate (L/h) / volume (L)).

| **D<sub>r</sub>** | **OD<sub>600</sub>** | **Ethanol** |  |  |  | **Glycerol** |  |  |  | **Acetic Acid** |  |  |  |
| --- | --- | --- | --- | --- | --- | --- | --- | --- | --- | --- | --- | --- | --- |
|  |  | **g/L** | **g/(L·OD)** | **g/(L·h)** | **Qp/Qs** | **g/L** | **g/(L·OD)** | **g/(L·h)** | **Qp/Qs** | **g/L** | **g/(L·OD)** | **g/(L·h)** | **Qp/Qs** |
| **0.025** | 7.2 | 8.37 | 1.16 | 0.21 | 0.42 | 0.46 | 0.06 | 0.01 | 0.02 | 0.07 | 0.01 | 0.00 | 0.00 |
| **0.05** | 5.4 | 7.15 | 1.32 | 0.36 | 0.36 | 0.81 | 0.15 | 0.04 | 0.04 | 0.08 | 0.01 | 0.00 | 0.00 |
| **0.1** | 2.4 | 5.24 | 2.18 | 0.52 | 0.43 | 0.83 | 0.35 | 0.08 | 0.07 | 1.42 | 0.59 | 0.14 | 0.12 |

**Figure 2.**
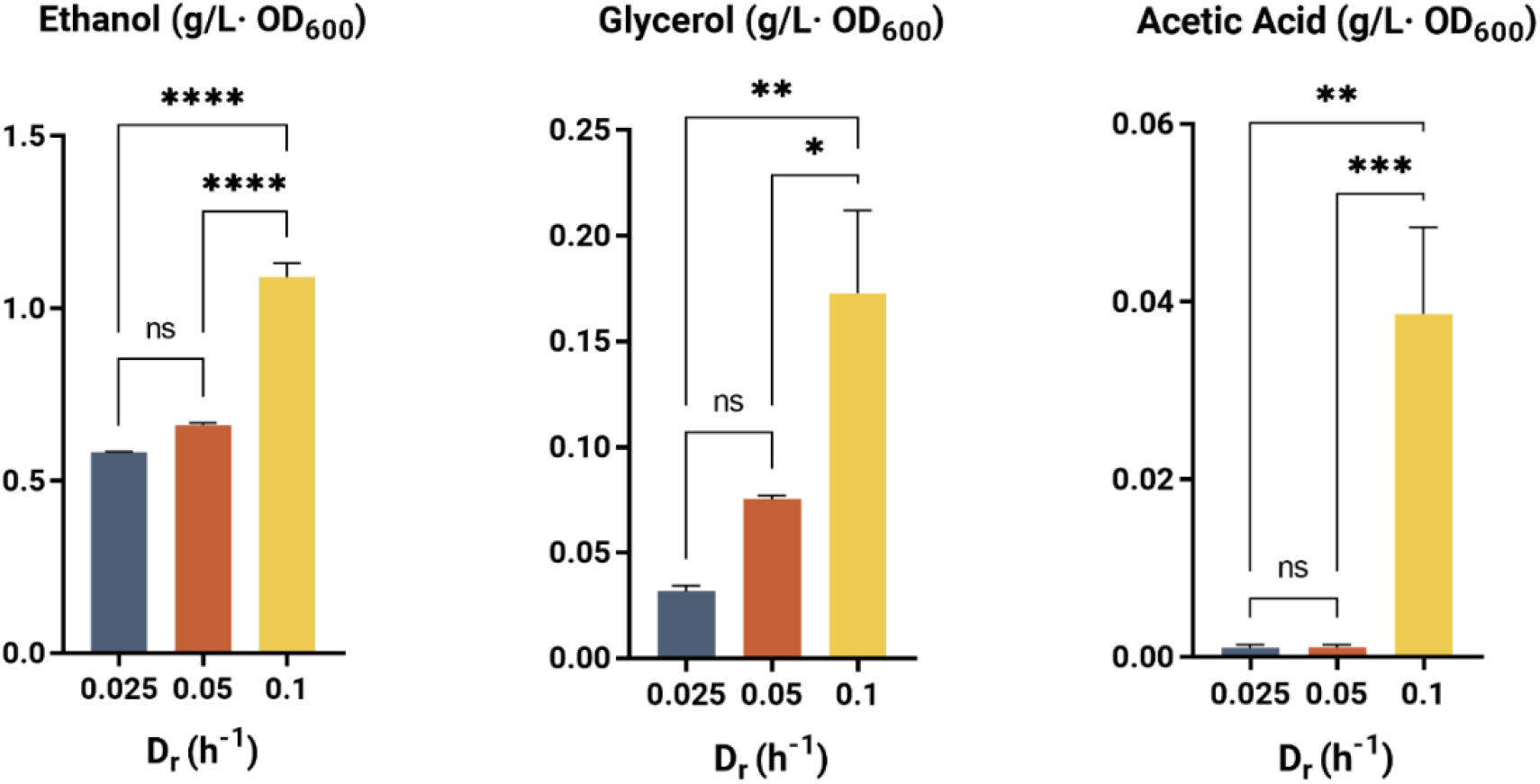
Effect of growth rate on metabolite production of NYCY 505 in synthetic medium. Specific concentrations of ethanol, glycerol, and acetic acid were measured by HPLC from steady-state samples of chemostats at different dilution rates and normalized by cell density as described Materials & Methods. Error bars represent the standard deviation of the mean of 3 consecutive steady-state samples of independent technical replicates. Statistical tests between sets were analysed using ANOVA with * p < 0.05 ** p < 0.01 *** p < 0.001 **** p <0.0001; ns =no significant change.

### Production of volatile compounds by NCYC 505 is affected by growth rate

The efflux samples of steady state chemostats operated at different dilution rates were assayed through SPME GC-MS for the most abundant volatile compounds present in alcoholic beverages. To be able to establish a clear, reproducible, and unambiguous correlation between growth rate and aroma compound production, a standard synthetic medium was utilized. Six compounds, belonging to three different classes; aldehydes, esters, and higher alcohols, and relevant to the beverage industry, were successfully semi-quantified.

Firstly, we found that the specific acetaldehyde concentration followed the trend of its oxidation product, acetic acid, detected previously via HPLC (Figure 3). A spike in acetaldehyde production at 0.1 h^−1^ D_r_ was also observed, which can be ascribed to an effect of overflow metabolism and an oversaturated flux towards ethanol fermentation. Acetaldehyde is the most abundant aldehyde in beer (23) contributing a pleasant, fruity aroma at concentrations lower than its odour threshold (10ug/g) (24), thus is an important metabolite to monitor.

**Figure 3.**
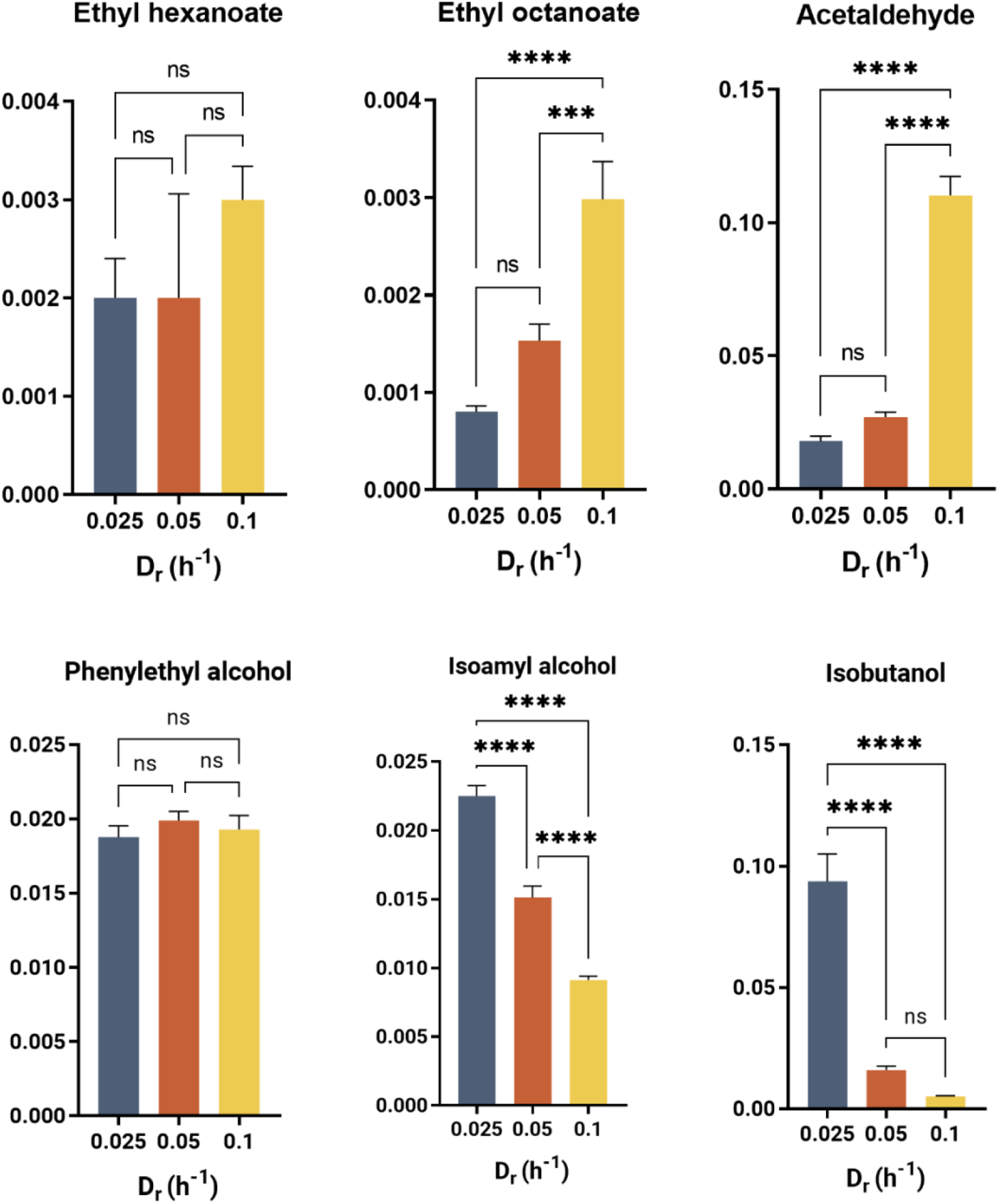
Volatile compounds production is greatly affected by growth rate in chemostats cultivation of NYCY 505 in synthetic media. Semi quantification measurement of volatile compounds were obtained through GC-MS-SPME. Error bars represent the standard deviation of the mean of 3 consecutive steady-state samples of independent technical replicates. Statistical tests between sets were analysed using ANOVA with * p < 0.05 ** p < 0.01 *** p < 0.001 **** p <0.0001; ns =no significant change.

Ester production was similarly affected by the growth rate. There was a positive linear correlation between growth rate and ethyl octanoate production, the most abundant ethyl ester in conventional brewing fermentations (25). Moreover, an increase in the specific concentration of ethyl hexanoate was recorded from 0.25 to 0.1 h^−1^ D_r_ (Figure 3). This finding suggest that there is a link between ester production and yeast growth. Another study also reported an increasing production of ethyl esters during the log phase, which decreased gradually after the consumption of 70% of the sugar content (26). Moreover, as ethyl ester formation is dependent on the concentration of its precursors, ethanol and acyl-CoA, this result is consistent with the higher specific concentration of ethanol we detected at increasing dilution rate.

Higher alcohols production, conversely, mainly decreased with dilution rate with both isobutanol and isoamyl alcohol exhibiting a clear downward trend, while phenylethyl alcohol specific concentrations remained stable across the different dilution rates (Figure 3). This result underlines the association between production of higher alcohols and cell growth. Although this link has already been suggested in literature (27, 28), as higher alcohol production results from amino acid metabolism via the Ehrlich pathway, our study further show that its production is also dependant on the physiological state of the cell, and that it is enhanced when the growth rate is severely limited by the carbon source. The observed trend of higher alcohol production is opposite to that of glycerol at higher dilution rates. Thus, while higher alcohols are thought to have a possible role in the maintenance of the redox balance (Quain, 1985), in the *S. cerevisiae* strain NCYC 505, glycerol production seems to be the preferred and predominant pathway to regenerate NAD^+^. Previous works primarily focused on the modulation of higher alcohol levels through an increase in the nitrogen and amino acid content in the medium (30, 31). Our results suggest instead that a modulation of the carbon source by the use of a defined feed profile could improve their titres, similar to the processes for the production amino acids (32, 33). Since higher alcohol production is dependent on the availability of their respective amino acid precursor (Figure 4A), the intracellular levels of phenylalanine, leucine and valine were quantified, being the precursors of phenylethyl alcohol, isoamyl alcohol and isobutanol, respectively. Leucine and valine showed a significant decrease in their specific concentration at 0.05 and 0.1 h^−1^ D_r_ (Figure 4B), consistent with the downward trend in the production of their respective higher alcohols observed at high dilution rate. Phenylalanine showed no significant difference in intracellular levels between the chemostats operated at 0.025 and 0.05 0.1 h^−1^ D_r_. However, the decrease recorded at 0.1 0.1 h^−1^ D_r_, while comparable with the one observed in leucine and valine levels, did not impact the specific concentrations of phenylethyl alcohol produced. Thus, the decreasing availability of intracellular amino acids may not be the only factor in play in the regulation of this pathway at different dilution rates.

**Figure 4.**
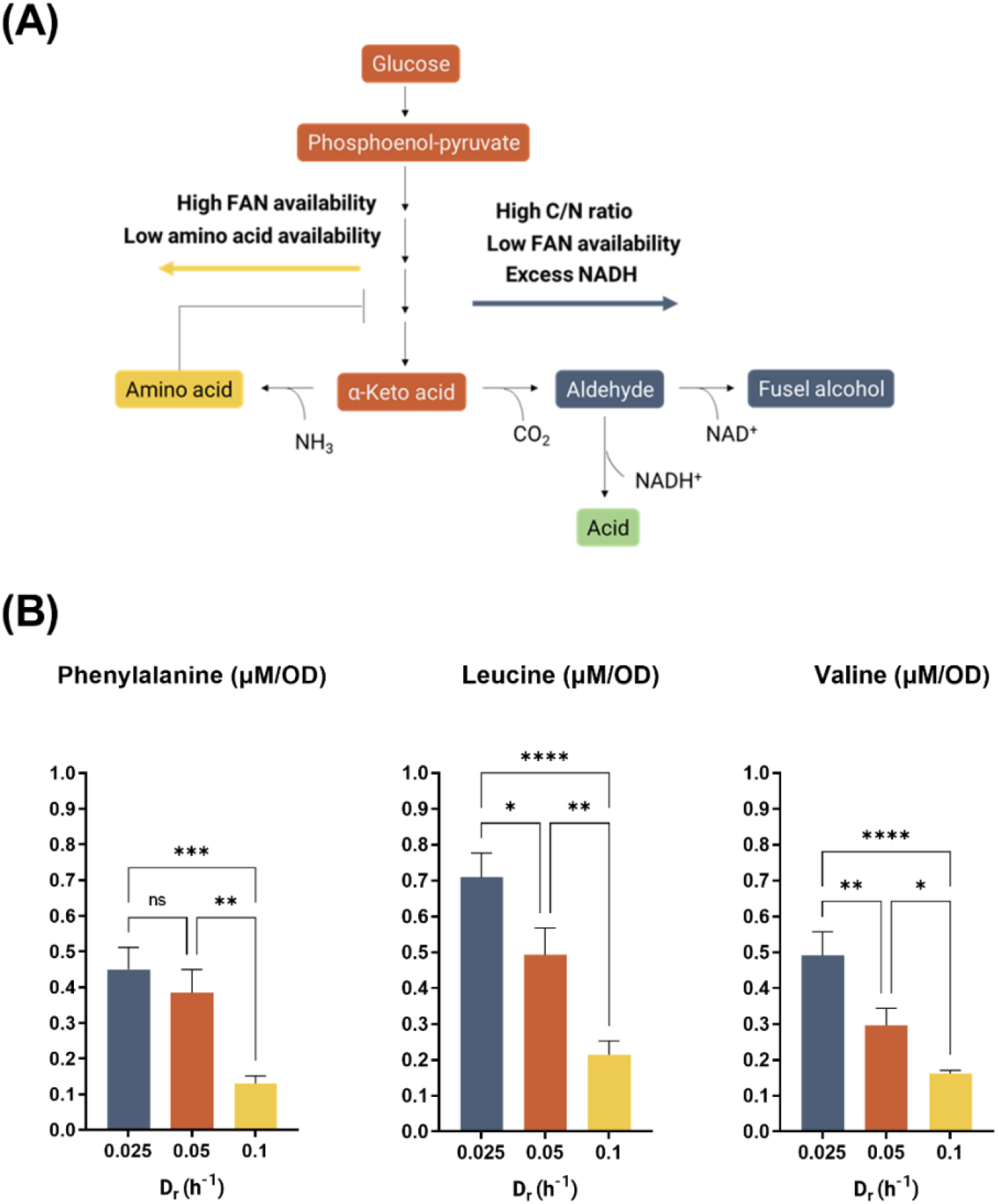
**(A) Anabolic pathway of higher alcohol production.** An α-Keto acid, derived by the glycolytic pathway or by the combination of glycolysis and the pentose-phosphate pathway can be transaminated to produce the corresponding amino acid or carboxylated to an aldehyde. The metabolic fate of the α-Keto acid is regulated by free amino nitrogen (FAN) availability, the concentration of amino acid, which regulates the pathway through a negative feedback loop, and by the carbon to nitrogen (C/N) ratio (26, 34). **(B) Intracellular amino acid specific concentrations.** Quantification of intracellular amino acids was obtained through LC-MS/MS. Error bars represent the standard deviation of the mean of 3 consecutive steady-state samples of independent technical replicates. Statistical tests between sets were analysed using ANOVA with * p < 0.05 ** p < 0.01 *** p < 0.001 **** p <0.0001; ns =no significant change.

### Controlled feeding profiles can be used to modulate the production of aroma compounds in conventional fermentations

To evaluate the effect of growth rate in the production of aroma compounds in conventional fermentations, the concentrations of selected compounds were assayed at different timepoints in batch and fed-batch experiments (Figure 1). NCYC 505 was grown in batch in 6.7 g/L YNB + 4% glucose at 20°C and in fed batch where the starting nutrient concentrations were halved compared to the batch and the remaining nutrients were fed at a constant flow over the course of 20h after 48h. Thus, the growth in the fed-batch experiment was artificially limited after 48h, to replicate a lower growth rate.

Profiles of growth, glucose consumption and ethanol productions are shown in Figure 5. Interestingly, no significant difference in ethanol and glycerol concentrations were found at the end of fermentations, with their concentration increasing linearly with glucose consumed.

**Figure 5.**
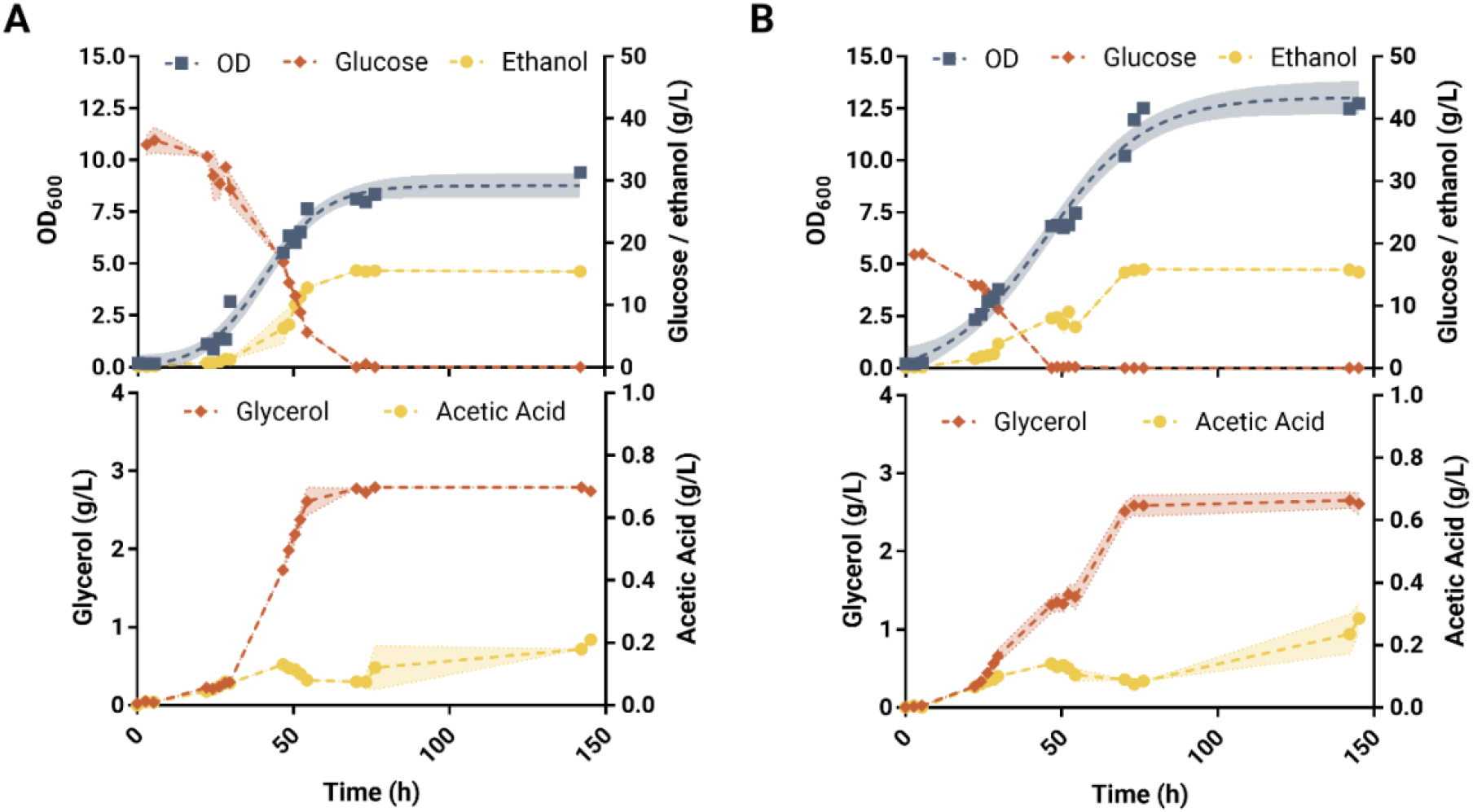
Profile of growth, glucose consumption and ethanol, glycerol, and acetic acid production in batch (A) and fed batch (B) fermentation of NCYC505 at 20°C. Standard deviation of two biological replicates is shown in lighter shades in the lines connecting each sample.

End-point fermentation samples were obtained after 76h of fermentation, after glucose was depleted in both batch and fed-batch fermentations, and after 146h. The end-point fermentation samples were assayed through SPME-GC-MS for esters and higher alcohol concentrations to evaluate their production profile and final titres (Figure 6).

**Figure 6.**
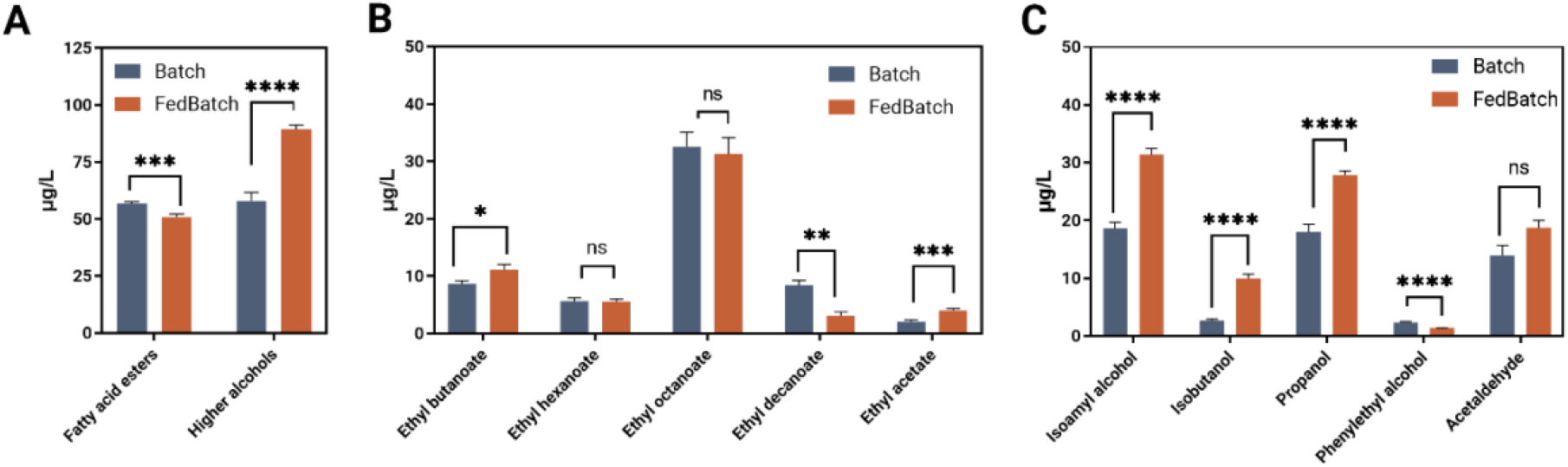
A controlled feeding profile can modulate the production of volatile compounds. **(A)** Cumulative concentrations of the detected fatty acid esters (ethyl butanoate, hexanoate, octanoate, caprinate) and higher alcohols (isoamyl alcohol, isobutanol, propanol, phenylethyl alcohol). **(B)** Quantitative measurement of esters, and **(C)** higher alcohols and acetaldehyde were obtained through GC-MS-SPME after 76h of fermentation in batch and fed batch. Error bars represent the standard error of biological replicates. Statistical tests between sets were analysed using t-test with * p < 0.05 ** p < 0.01 *** p < 0.001 **** p <0.0001; ns =no significant change.

Thanks to the higher concentrations reached in the end-point samples compared to the chemostat fermentations, it was possible to increase the number of volatile compounds assayed with five esters (ethyl butanoate, ethyl hexanoate, ethyl octanoate, ethyl decanoate and ethyl acetate), four higher alcohols (isoamyl alcohol, isobutanol, propanol and phenylethyl alcohol), and acetaldehyde being quantified. Moreover, the concentrations of these compounds were determined throughout the course of the fermentation to highlight the active phases of production (Figure S1).

The production of seven out of ten volatile compounds were found to be modulated by the feeding profile. The batch experiment reached a higher concentration of fatty acid esters and significantly lower titres of higher alcohols, confirming the general trends observed in chemostat (Figure 6A).

However, no significant difference was observed in the production of ethyl hexanoate and ethyl octanoate concentrations between the two experiments after 76h, which in chemostat experiments resulted enhanced by growth rate. Conversely, ethyl butanoate titres slightly increased in fed-batch by 2.45 μg/L, while ethyl decanoate production was almost tripled in batch. These results indicate that high growth and sugar concentrations mainly stimulate the production of esters derived from longer-chain fatty acids, as in the case of ethyl decanoate.

The prolonged period of growth in the fed-batch fermentation resulted in higher levels of ethyl acetate, an acetate ester produced by a condensation reaction between acetyl-CoA and ethanol (27) (Figure 6B). Production profile of ethyl acetate followed the one observed for ethanol in both experiments. Thus, the ethanol fermentation over the course of the entire 76h in the fed-batch might be a result of the higher concentrations of ethyl acetate, compared to the batch. Acetaldehyde titres increased the most in the first 30 hours of fermentation, both in batch and fed-batch (Figure S1). The concentration remained mostly stable during the course of the fermentation, as a result, the final concentration of acetaldehyde did not differ between batch and fed-batch fermentations (Figure 6C).

Consistent with the results observed in chemostat fermentations, a 60% increase in the levels of total higher alcohols was recorded in fed-batch fermentation, where the lower availability of glucose resulted in fewer hours of sustained specific growth rate compared to the batch (Figure 6C). In particular, the feeding protocol in fed-batch resulted in a 2.5-fold increase in the levels of propanol, and over 50% higher titres of isoamyl alcohol and isobutanol (Figure 6C). Phenylethyl alcohol levels resulted higher in batch as its production was greatly reduced in the second phase of the fed-batch fermentation (Figure S1). In the initial stages of the fermentation, the yield on glucose of phenylethyl alcohol was similar in both experiments (0.49 μg/g in batch and 0.47 μg/g in fed-batch). However, the yield in the fed-batch decreased dramatically to 0.22 μg/g when new nutrients were drip-fed into the media while the batch reached a final yield 0.60 μg/g. Thus, we propose that phenylethyl alcohol production in NCYC 505 might be enhanced when nutrients levels in the media are in excess, and decrease when nutrients become limiting, as observed in fed-batch fermentations.

Lower levels of higher alcohol were reported to be a cause of unbalanced flavour profile higher in high-gravity brewing, characterised by high starting sugar concentrations (35, 36). The result obtained suggests that the final titres of higher alcohols could be effectively modulated through the use of fed batch fermentations.

## Conclusions

The incredible recent growth of the craft beer market has fuelled the need to develop new beverages catering to the consumer need for a wider choice of flavours and beery style. While synthetic biology approaches helped identify and modulate the pathways responsible for aroma compound production, the limitations in the use of genetically modified organisms in food and beverages has restricted these applications. Recent developments in the field of yeast hybridisation have helped develop new strains able to produce new and complex aroma profiles (37–39).

However, little is known about the physiological role of aroma compound production in yeast, and this is hindering evolution experiments. Thus, it is of foremost important to dissect the pathways connected with aroma profile and understand how they can be manipulated to enhance the aroma profile of fermented beverages. In this study, a potential link between the production of aroma compounds and growth rate was investigated in the *Saccharomyces cerevisiae* type strain NCYC 505.

Continuous fermentation experiments in anaerobic conditions at different dilution rates allowed unambiguous correlation of yeast growth rate with the production of four out of the six volatile compounds assayed. In particular, ethyl octanoate and acetaldehyde levels increased with growth rate while the synthesis of two higher alcohols, propanol and isoamyl alcohol, was found to be inversely correlated with growth. The enhanced production of higher alcohols was also found to be directly correlated with the intracellular levels of their respective amino acid precursor, which were more abundant at lower dilution rates. Moreover, higher alcohols were found to play little to no role in redox balance at high dilution rate, with glycerol synthesis being the preferred pathway to regenerate NADH used in anabolic reactions.

The continuous fermentation experiment confirmed esters and higher alcohols as tightly linked with cell metabolism and suggest that their metabolism is influenced by the physiological state of the cell. Thus, modulation of the nutrient availability might be an important tool for exerting control on the final aroma profile of a fermented product. Guided by these results, fermentation experiments in batch and fed-batch were devised to investigate the extent the aroma profile could be manipulated by limiting nutrients availability. The concentrations of seven out of ten volatile compounds detected were found to be significantly different in the end-point sample of the fermentations. In particular, higher alcohols levels increased by 60% in fed-batch fermentations where yeast growth was limited by nutrients availability, with propanol increasing 2.5-fold over batch levels, while ethyl decanoate concentrations tripled in batch fermentation. Thus, the newly described link between growth rate and aroma compound production was successfully exploited to increase the titres of higher alcohol in NCYC 505 and carefully modulate its aroma profile through rational design of the fermentation process. The implementation of feeding profiles to artificially regulate growth will be an important tool to cater to consumer needs and generate new fermented beverages harbouring new and varied aroma profiles.

## Materials and Methods

### Strain and media

All experiments were performed with the industrial yeast *S. cerevisiae* type strain NCYC 505 purchased from the National Collection of Yeast Cultures (NCYC). The yeast was maintained on YPD agar (15 g/L agar, yeast extract, 10 g/L, peptone, 20 g/L, glucose, 20 g/L). Precultures were grown overnight in shake flask cultures on 50 ml YPD (yeast extract, 10 g/L, peptone, 20 g/L, glucose, 20 g/L) at 30°C. Biomass was recorded via optical density measurement at 600nm with an Eppendorf BioSpectrometer.

### Bioreactor fermentations

1L stirred tank bioreactors (Multifors, Infors-HT, Bottmingen, Switzerland) were used to perform all fermentation experiments. Cells were grown in 500 mL of YNB + 2% glucose after inoculation with a washed overnight culture to reach an initial OD of 0.1. Temperature was controlled at 20°C and agitation set at 300 rpm. pH was adjusted by the addition of 2M sodium hydroxide (NaOH) to maintain a constant value of 5. An influx of oxygen was maintained for 5 min to saturate the vessel with air, prior the start of the fermentation. After inoculation, oxygenation was set to 0%. Aliquots were collected at different time intervals for optical measurements reading. The samples were then centrifuged (13,000 g for 5 minutes) and stored at −20°C for further analysis.

Fed-batch fermentations were performed as described above in 500 ml of 3.35 g/L YNB + 2% glucose. After 48 h, 25 ml of 20% glucose (w/v) and 3,35 g/L of YNB were fed at a constant flow of 1 1.25 ml/·h for 20 h until the fermentation was reverted to batch.

### Continuous fermentations

For chemostats, the fermentation was started as described above in batch for the first 36h and then maintained in continuous for 10 days. A constant feed of fresh media was pumped in the fermenter at different dilution rate (D = 0.025, 0.05, 0.1 h^−1^) with a peristaltic pump (Infors-HT). The efflux was removed from a sampling tube connected to a peristaltic pump in order to keep the volume constant. The steady state was defined as the situation in which at least a volume change had passed after the last change in OD_600_ measured and where the measured ethanol and glycerol concentrations were constant.

### HPLC analysis

The concentration of metabolites (glucose, glycerol, acetic acid, and ethanol) was measured by HPLC using a 1260 Infinity II LC System with a Refractive Index Detector (Agilent). A 300×7.8 mm Hi-Plex HPLC Column (Agilent) was equilibrated with 5 mM H_2_SO_4_ in HPLC grade water at 55° at a 0.8 mL/min flow rate. Compounds were identified by retention times and quantified using calibration curves generated from authentic standards. For chemostat efflux samples the concentration is expressed as specific concentration, calculated dividing the concentration in g/L by the OD_600_ value for each timepoint.

### Aroma compound analysis

Volatile compounds were detected and analyzed by using a Thermo Scientific TSQ Quantum GC Triple Quadropole GC-MS. 0.5 ml of sample were prepared in a 20 ml vial added with 25 μl of the internal standard 2-octanol (2.5 mg/L) and 0.5 g of NaCl. The samples were incubated for 10 min at 40°C. The volatile compounds were collected on a Divinylbenzene–Carboxen- Polydimethylsiloxane 2 cm fiber (DVB-CAR-PDMS Supelco) for an extraction time of 30 min. A VF-wax column (Agilent) 30 m/I.D 0.25 mm/Film 0.25 μm was used for the separation.

The oven was kept at 40 °C for 4 min then increased by 6 °C/min to 250 °C and kept at the final temperature for 5 min. The injector and interface temperatures were kept at 250 °C as well. Helium was used as the carrier gas with a flow rate of 1.2 ml/min. The time for thermal desorption of analytes was 4 min. The MS detector was operated in full scan mode at 70 eV with a scan range from 40 to 300 m/z for 44 min.

Data analysis was performed using the software ThermoXcalibur (Version 2.2 SP1.48, Thermo scientific). Chromatogram peak annotation was based on the retention time of the reference standard and comparison with a mass spectral database (NIST version 2.0). The absolute concentration of the compound in the sample was calculated with calibration curves generated from pure reference standards. Measurement bias was corrected with the internal standard.

### Amino acid analysis

Intracellular amino acids were extracted as described in Gent and Slaugther (1998) (40). LC-MS/MS analysis was conducted using an ultra-performance liquid chromatography system (Waters Acquity UPLC H-class) coupled to a Xevo TQ-S triple-quadrupole mass spectrometer (Waters Corporation, MA, USA) equipped with an electrospray ionization source (ESI). The desolvation gas flow rate was set to 300L/h at a temperature of 600 °C. The cone gas flow rate was fixed at 150 L/h and the source temperature at 150 °C. The source offset was set to 50 V. The capillary voltage was optimized at 1.5 kV for negative mode (ESI-). The following compounds were monitored in ESI- mode as follows; Valine (SIR of mass 116.07, 0.2-1.2 min); Leucine/Isoleucine (SIR of mass 130.09, 0.75-1.4 min); Phenylalanine (MRM 164.07->146.60, 2.0-3.0 min). MassLynx v 4.1 (Waters) software was used to process the quantitative data obtained from calibration standards and from diluted samples. The mobile phases, columns, column temperature and gradient programs were set based on the specific chemical properties of the target analytes. A Waters Acquity BEH C18 column (50 mm × 2.1 mm, 1.7 μm) was used at 45 °C, flow rate 0.3 mL/min. An optimum separation gradient was obtained with a binary mobile phase A (H_2_O + 0.1% Formic Acid) and B (Acetonitrile +0.1% Formic Acid). The gradient elution program was: 0- 1.2 min at 100% A; 1.2 −2.5 min to 90% A, hold at 90% A for 2.5 min; reduce to 5% A over 0.5 min, hold for 0.5 min before returning to 100% A for 1 min. The total run time was 7 min. The inject volume was 5 μL.

## Acknowledgments

This work was supported by the European Commission (H2020-MSCA-ITN-2017) grant number 764364 awarded to DD and UV, and by Biotechnology and Biological Sciences Research Council (BBSRC) grant number BB/R013497/1. The authors would like to thank Clare Lin Lin for useful discussion on amino acids quantification. The authors also acknowledge the support received from the Manchester Institute of Biotechnology analytical facility and from the mass spectrometry facilities.

---

**Figure S1.**
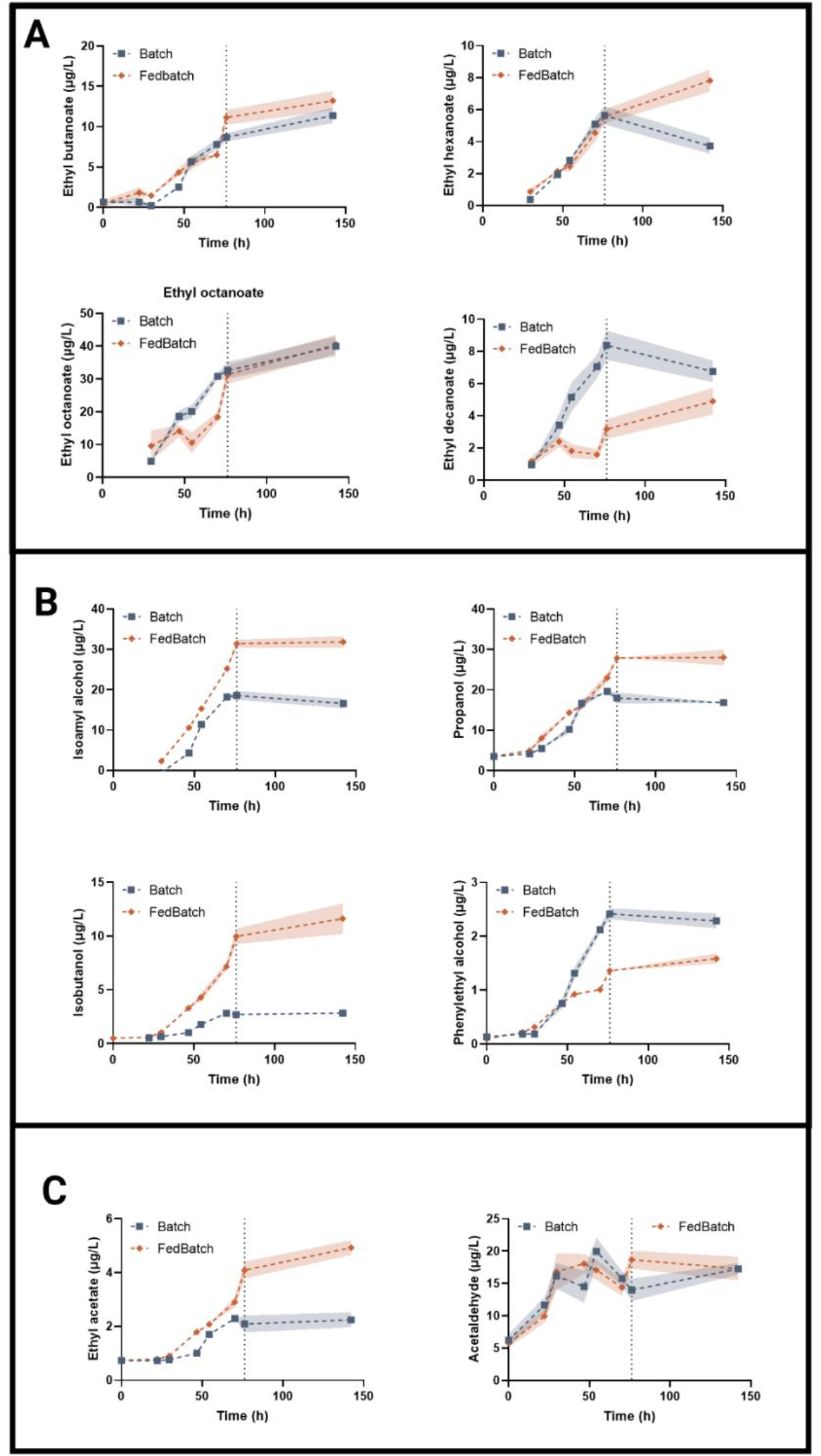
Production kinetics of (A) fatty acid esters, (B) higher alcohols, (C) ethyl acetate and acetaldehyde in batch and fed-batch fermentations. Standard deviation of two biological replicates is shown in lighter shades in the lines connecting each sample. A vertical dashed line represents the timepoint when glucose was depleted in both batch and fed-batch fermentations, after 76 hours.

